# Development of a compact alkynyl-enrichable crosslinker for in-depth in-vivo crosslinking analysis

**DOI:** 10.1101/2021.07.30.454285

**Authors:** Hang Gao, Lili Zhao, Baofeng Zhao, Zhou Gong, Qun Zhao, Lihua Zhang

## Abstract

Chemical crosslinking coupled with mass spectrometry (CXMS) has emerged as a powerfμl technique to capture the dynamic information of protein complexes with high sensitivity, throughput and sample universality. To advance the study of in-vivo protein structures and protein-protein interactions on the large scale, a new alkynyl-enrichable crosslinker was developed with high efficiency of membrane penetration, reactivity and enrichment. The crosslinker was successfully used for in-vivo crosslinking of intact human cells, resμlting in 6820 non-redundant crosslinks identified at a false discovery rate (FDR) of 1% using pLink 2.0, which 4898 (71.8%) of the cross-links were assigned as intraprotein and 1922 (28.2%) were interprotein links. To our knowledge, this is also the first time to realize the in-vivo crosslinking with a non-cleavable crosslinker for homo species cells.

## Introduction

In living cells, proteins interact with other proteins to form large complexes, which regμlate various life processes in a precise and orderly manner. Protein complexes perform specific functions by regμlating the dynamic changes of their conformation and interactions to form the high-dimensional signal transduction networks^[1-4]^. In constrast to in-vitro extracellμlar environment, the living cell system is characterized by high confined and crowding effect, making it with a unique subcellular space resolution and strong viscosity features to exactly maintain the protein complexes’ natural conformation and dymanic interaction assemby for function execution. Thus, efficiently and credibly characterizing the conformation and interactions of protein complexes in living cells is essential for comprehensive understanding the functional mechanism of complicated biological processes.

Recently, chemical crosslinking coupled with mass spectrometry (CXMS) has emerged as a powerful technique to obtain the dynamic structural information and direct interactions of protein complexes, due to its advantages of high sensitivity, throughput and sample universality without any requirement on the protein molecular weight and purity^[5-9]^. Typically, complemented to existing techniques for static protein complex analysis, such as X-ray, Cryo-EM, AP-MS, et al., CXMS can also analyze the dynamic conformation and interactions of protein complexes by providing the ensemble distance constraints of structural sites within the time limit of crosslinking. In CXMS technology, it utilizes special small molecular reagents (namely crosslinkers) containing two reactive groups separated by a spacer to bridge two amine acid residues in close proximity, followed by shot-gun based crosslinked protein digestion and LC-MS analysis for crosslinks identification^[10-12]^.

Continued advances made in recent years to various crosslinkers, crosslinking methodology, mass spectrometry instrumentation and high-end data processing softwares enable CXMS to be applied to diverse biological systems, especially in living systems^[13-16]^. To realize the in-vivo crosslinking, formaldehyde was a promising crosslinker due to its small size, high reactivity and fast cell permeability. However, the broad specificity and chemical diversity make identification of the end-product difficult, particularly from very complex biological samples, limiting the application of formaldehyde^[2,3]^. In recent years, in-vivo large-scale analysis of cellular interactome dynamics have been reported, such as using an enrichable protein interaction reporter (PIR) crosslinkers, observing the changes of protein conformations and interactions during paclitaxel treatment^[17]^. For in vivo sub-organelle chemical crosslinking, DSSO was successfully investigated the organization of proteins in native condition, providing direct evidence for oxidative phosphorylation comolexes assembly in mitochondria^[10]^. These in-vivo advances greatly increase our ability to discover the structural dynamics of intracellular protein complexes, leading to discovery of the diverse functional regulation related to specific structural basis, which is extremely difficult to achieve in vitro.

Identification coverage of CXMS directly determines its ability for protein complex analysis. However, low-abundance cross-linked peptides of interested were produced, due to the limited reaction efficiency, relatively few amino acids exposed to protein surface, and the diversity of crosslinking products. The difficulty of identifying these low-abundance peptides is compounded by the increasing complexity of the biological samples, especially for in-vivo crosslinking with high matrix background interference. As a response, many strategies have been developed to enhance the enrichment of cross-linked peptides, including chromatographic techniques such as strong cation exchange (SCX) or size exclusion chromatography (SEC) or directly utilization of enrichable trifunctional crosslinkers^[2-4]^. Among them, the development of enrichable crosslinkers was the most promising strategy for crosslinked peptides enrichment, such as Leiker^[18]^, Phox^[19]^, Azide-A-DSBSO^[20]^, BDP-NHP^[21]^, et al., exhibiting a high enrichment efficiency and specificity in complex biological samples. It’s worth noting that the extraordinary binding strength and highly specific biotin−streptavidin system makes biotin moiety widely used in protein labeling and enrichment^[18, 21, 22, 23]^. However, these crosslinkers integrated with bulky biotin are not ideal for in vivo crosslinking, due to the steric hindrance and strong hydrophobicity. Instead, the biotin enrichment handle can be indirectly functionalized transformations utilizing smaller azide or alkyne that allow performing copper-catalyzed azide-alkyne 1,3 dipolar cycloadditions (CuAAC) after crosslinking reaction^[23]^. The prominent advantages of this approach are the complete orthogonality in biological systems, great specificity and sensitivity without significant competing reactions, and better biological compatibility in mild aqueous conditions.

Inspired by the great progress made in achievement of in-vivo crosslinking and efficient cross-links enrichment, a new compact membrane-permeable crosslinker was developed, possessing trifunctional groups of two widely used *N*-hydroxysuccinimide (NHS) esters with high reactivity targeting to the ε-amino group of lysine side chain under physiological condition, and an alkynyl enrichment group to enable sensitive cross-links enrichment, termed bis-succinimidyl bearing propargyl tag (BSP). With this crosslinker, intracellular crosslinking in living cells can be achieved, followed by a sandwich two-step enrichment method based on click chemistry-affinity enrichment. Finally, the method was successfully used for in-vivo crosslinking of protein complexes in intact human cells. To our knowledge, this is also the first time to realize the in-vivo crosslinking with a non-cleavable cross-linker for homo species cells.

## Experimental Procedures

### Materials and Reagents

The chemical reagents were obtained commercially and used as received without further purification. Synthetic peptide Ac-SAKAYEHR (99.14% purity) was custom ordered from ChinaPeptides (Shanghai, China). Bovine serum albumin (≥98% purity) and diazo biotin−alkyne were purchased from Sigma-Aldrich. Streptavidin agrose resin was purchased from Thermo Fisher.

### Crosslinking of Standard Peptide

Synthetic peptide Ac-SAKAYEHR was first dissolved in 100 μl anhydrous dimethyl sulfoxide (DMSO) containing 1% triethylamine (v/v). BSP was reconstituted in anhydrous DMSO and mixed with peptide solution in molar ratio of 1:2. After a quick vortex, the crosslinking reaction was performed at room temperature for 5 min. Adding 20 volumes of 0.1% TAF/H_2_O to quench the reaction. DMSO was removed by C18 tips (similar to Peptide Desalting), followed by drying on a speed-vac concentrator. The resulting crosslinked peptides were diluted to 1 μg/μl in 0.1% FA/H_2_O prior to MALDI-TOF MS and ESI-MS analysis.

### Crosslinking of BSA for SDS-PAGE analysis and Mass Spectrometry Acquisition

Crosslinking of BSA for SDS-PAGE analysis. The BSP and DSS (Thermo Fischer) were freshly dissolved in anhydrous DMSO with a concentration of 500 mM and 50 mM, respectively. BSA was dissolved in PBS (pH 7.4, Thermo Fischer) with a concentration of 1 mM. The BSA concentration was diluted with PBS and mixed with BSP and DSS in different molar ratio from 1:5 to 1:50. The reaction mixtures were incubated at 25 °C for 5min with soft agitation, followed by quenched with 50 mM ammonium bicarbonate (ABC) solution. After addition of 5-fold excess acetone (pre-cooling to -20 °C), the proteins were precipitated at -20 °C overnight to remove the non-reacted crosslinkers. The samples were centrifuged at 4000g for 5 min and washed once with cold acetone. The resulting proteins were resuspended in 1х loading buffer (Bio-Rad) and boiled at 95 °C for 5 min. A separation gel of 12% loaded with 10 µg protein for each aisle was run at 120 V for 90 min. After fixed in 50% MeOH and 10% AcOH for 30 min, the gel was stained with coomassie brilliant blue for one hour and destained in water overnight. The picture was taken by bio-rad molecular imager and handled with Image Lab software.

Crosslinking of BSA for mass spectrometry acquisition. A volume of 100 μL of 100 μM BSA in PBS was incubated with freshly dissolved BSP at a molar ratio of 1:50 for 5 min followed by addition of 50 mM ABC to quench the crosslinking reaction. After acetone precipitation at -20 °C overnight, the pellet was obtained by centrifugation (4000 g, 5 min, 4 °C). The protein pellet was redissolved in 8 M urea and reduced with 10 mM TCEP for 2 h at room temperature, followed by alkylation with 20 mM IAA for 30 min at room temperature in the dark. The solution was diluted by 7-fold volume 50 mM ABC to reduce the urea concentration to less than 1M. The sample was digested with trypsin at an enzyme-to-protein ration of 1:50 (w/w) at 37 °C for 8 h after which 1% formic acid (v/v) was added to quench the digestion. After desalted with C18 tips, the digestion was dissolved in 0.1% SDS/PBS (v/v) and subjected to copper-catalyzed click chemistry reaction.

### Click chemistry reaction to introduce a biotin handle

The click chemistry reaction was carried out by addition of 1 mm diazo biotin−alkyne (DBA), 4 mm sodium ascrobate, 4 mm THPTA and 4 mm CuSO_4_ at 37 °C for 3 h. After reaction, the excess reagents of sodium ascrobate, THPTA and CuSO_4_ were removed by C18 tips and the excess DBA probe was removed by MCX column (Waters, Oasis, Made in Ireland) that may potentially interfere with the subsequent enrichment.

### Enrichment and release of Biotinylated Peptides for Mass Spectrometry Acquisition

The streptavidin agrose resin was firstly washed by 50 mM ABC for three times before use. The biotinylated peptide samples were redissolved in 50 mM ABC, mixed with appropriate amount of streptavidin agrose resin (make sure the supernatant was colourless) and slightly agitated at room temperature for 2 h. The supernatant was removed completely by centrifuging at 2000 g for 5 min, and then the streptavidin beads was washed with 1 M KCl, H_2_O, 10% CH_3_CN/H_2_O (v/v), H_2_O in turn for two times. After extensive washing, the peptides captured by streptavidin beads were cleaved by one column volume elution buffer (300 mM Na_2_S_2_O_4_, 6 M urea and 2 M thiourea dissolved in 20 mM HEPES, pH 8.0) at 37 °C for 30 min and collected by centrifugation. Repeat the elution procedure to ensure complete cleavage and peptide recovery. Combine the elution supernatant and the peptides were desalted by C18 tips, resolubilized in 0.1% FA/H_2_O (v/v) followed by mass spectrometry acquisition.

### Fluorescent confocal microscopy of BSP cell membrane permeability

The cell membrane permeability of BSP was studied by fluorescent confocal microscopy. For confocal microscopy samples, Bel7402 were grown under a humidified atmosphere containing 5% CO_2_ at 37 °C in RPMI 1640 Medium containing 10% fetal bovine serum and 1% penicillin/streptomycin in 35-mm Petri dishes with number 1.5 cover glass bottom for two days until they reached 70-80% confluence. The adherent cells were washed five times with PBS and then reacted with 5 mM BSP in 1 ml PBS containing 1% DMSO (v/v) at room temperature for 0.5 min, 1 min, 3min, 5 min, respectively. Following the crosslinking reaction, the cells were again washed 5 times and fixed by 1 ml 10% HCHO/PBS for 15 min at room temperature with constant shaking. After fixation, the cells were incubated with 0.1% triton X-100 (v/v) in 1ml PBS for 60 min for cell membrane perforation. The click chemistry reaction was processed with 1ug/ml FITC-N_3_ (Sigma-Aldrich) in the mixture of 4 mm sodium ascrobate, 4 mm THPTA ligand and 4 mm CuSO4 for 3 h at room temperature in the dark. After extensive washing with PBS, the cells were incubated with 1 ug/ml DAPI for 15 min for cell recognition and localization. The confocal microscopy samples were finally prepared after washing the cells with PBS for three times. The blank control was carried out in 1% DMSO/PBS for 5 min which was the only difference from the experimental groups. Confocal fluorescent imaging was performed separately in 405 nm and 488 nm channels using a 100X objective by andor live cell confocal imaging platform (Nikon Instruments Inc., USA).

### Preparation of In-vivo crosslinking of Bel7402 cells

Bel7402 cells were grown as described above in 15-cm plates until they reached 80% confluence. Cells were harvested by directly scrape and collected into centrifuge tubes. The cells were pelleted by centrifuging at 500 g for 5 min at 4 °C and washed 3 times with PBS. Cell counting was performed before the crosslinking reaction to roughly estimate protein concentration. Resuspending the cell pellet into the freshly dissolved BSP at a concentration of 5 mM in 1% DMSO/PBS and the crosslinking reaction was carried out at room temperature for 5 min with slightly agitation followed by washing 3 times with PBS. After centrifugation, the cell pellet was resuspended in 1% SDS/PBS containing 1% protease inhibitor cocktail (v/v) and lysed by sonication at amplitude 50% power in ice (5 s on, 5 s off) until the sample is no longer viscous. The following steps (acetone precipitation, reduction, alkylation, protein digestion, click chemistry procedure, enrichment and release of biotinylated peptides) were similar to that of BSA samples.

### Peptide samples fractionation by high pH reverse-phase C18 column

After eluted from streptavidin resin, the crosslinked peptides obtained from in vivo crosslinked cells were fractionated by high pH reverse-phase C18 column (Durashell, 5 um, 100 Å, 2.3 × 150 mm i.d.). The fractionation solvent A consisted of 2% CH_3_CN in 98 % H_2_O, solvent B consisted of 2% H_2_O in 98 % CH_3_CN, and both were adjusted to pH 10 by ammonium hydroxide. The fractionation gradient was set as follows: 0-10 min (0 B, desalting step), 10-10.1 min (2% B), 10.1-45 min (2-30% B), 45-60 min (30-45% B), 60-70 min (45-90% B), 70-80 min (90% B). We collected 20 fractions and dried followed by LC-MS acquisition.

### Mass Spectrometry Acquisition

#### MALDI-TOF analysis of standard crosslinked peptides

The resulting crosslinked samples from standard peptide (crosslinked sample before click chemistry, biotinylated sample after click chemistry reaction and elution sample after enrichment and cleavage from streptavidin beads) were diluted in 0.1% FA/H_2_O. The the samples were spotted onto the MALDI plate and mixed with 2,5-Dihydroxybenzoic acid (DHB, as a matrix). Mass spectra were manually acquired in a positive linear mode by using a ultrafleXtreme MALDI-TOF mass spectrometer (Bruker Daltonics, Germany) and the FlexControl software (version 3.3). The instrument parameters were set on the FlexControl software (positive linear mode; laser frequency 200 Hz; 800-3000 KDa molecular weight range; 20.0x detector gain). Each mass spectrum was generated from several single laser shots in 200-shot from different positions of the sample to generate the spectra with an intensity ≥10^4^ arbitrary units.

#### Orbitrap Flusion Lumos acquisition of cross-linked peptides

The desalted or fractionated samples resolubilized in 0.1% FA/H_2_O (v/v) were centrifuged at 16000 g for 30 min and the concentration of peptides were measured by Nanodrop one Microvolume UV-Vis Spectrophotometer (Thermo Fisher). All peptide samples were analyzed in technical triplicate by LC-MS using an Easy-nano LC 1200 system coupled to an Orbitrap Fusion Lumos mass spectrometer (Thermo Fisher). The samples were automatically loaded onto a C18 trap column (150 µm i.d. х 3 cm) and separated by a C18 capillary column (150 µm i.d. х 15 cm), packed in-house with ReproSil-Pur C18-AQ particles (1.9 µm, 120 Å). Mobile phase solvent A consisted of 0.1% FA in HPLC H_2_O, and mobile phase solvent B consisted of 0.1% FA in 80% acetonitrile and 20% HPLC H_2_O. The unfractionated cell samples were separated using a 150 min gradient as follows: 53 min from 8% to 20% B, 50 min from 20% to 35% B, 25 min from 35% to 50% B, 2 min from 50% to 95% B and 95% B maintained for 20 min. The crosslinked standard peptides, BSA digestion and fractionated intact cell samples were separated using a 85 min gradient as follows: 15 min from 10% to 20% B, 35 min from 20% to 35% B, 25 min from 35% to 50% B, 1 min from 50% to 95% B and 95% B maintained for 9 min. The mass spectrometry was operated in data-dependent acquisition mode with one fμll MS scan 350-1500 at R = 60, 000 (m/z = 200), followed by MS/MS scans at R = 15, 000 (m/z = 200), RF Lens (%) = 30, with an isolation width of 1.6 m/z. The AGC target for the MS1 and MS2 scan were 400000 and 50000, respectively, and the maximum injection time for MS^1^ and MS^2^ were 50 ms and 30 ms. The precursors with charge states 3 to 7 with an intensity higher than 20, 000 were selected for HCD fragmentation, and the dynamic exclusion was set to 20 s. The normalized collision energy was set as 30% (HCD).

#### Data Analysis

The raw MS data were analyzed by pLink 2.0 software (version 2.3.5) to identify the crosslinking information. The search parameters were set as follows: precursor mass tolerance 20 ppm, fragment mass tolerance 20 ppm, precursor filter tolerance 10 ppm, crosslinker BSP (cross-linking sites K and protein *N*-terminus, cross-link mass shift 411.191, monolink mass shift 429.201), trypsin digestion up to 3 missed cleavages, peptide mass from 500 Da to 6000 Da, peptide length from 5 to 60, fixed modification carbamidomethyl [C], variable modification oxidation [M] and acetyl [Protein N-term]. For BSA samples, the BSA sequence downloaded from UniProt was used as database with a separate FDR ≤ 0.01 at PSM level and searching resμlts were manually filtered by E-value ≤ 0.001, PSM ≥ 4 for the identified residue pairs. For intact cell crosslinking samples, the data were searched against UniprotKB human protein database from April 2019 containing 42432 proteins. The FDR was set to 0.01 at PSM level to control the data threshold separately.

## Results and discussion

### Design of BSP to enable in-vivo crosslinking

To maintain the natural conformation of the protein complexes in living cells, crosslinking reaction occurred in the intracellular aqueous environment was required. However, due to the demands for multifunctional groups and membrane lipid solubility, the existing membrane-permeable crosslinkers often have relatively strong hydrophobicity. Thus a high proportion of organic solvents, such as DMSO, was usually needed to assist the crosslinkers dissolution, which might disturb the structure of protein complexes to some extent, resulting in artificial false positive results. Hence, the design of a crosslinker with better solubility will be more suitable for in-vivo crosslinking. Besides, a compact size was required for crosslinker to enable higher cell membrane permeability and lower reactive sterical hindrane, which will be beneficial to capture the dynamics of protein complexes in living cells.

Based on the above, an enrichable trifunctional crosslinker, termed as BSP, was designed (Figure 1A) and synthesized (detailed in supplementary material) consisting of two lysine-targeting reactive groups N-hydroxysuccinimide (NHS) esters and one orthogonal click-enrichable group alkyne, with maximum Cα–Cα distance restraint of 27.9 Å (Figure 1B). As comparison, the amphipathy and molecular size of BSP were calculated with the reported chemical crosslinkers, as shown in the Figure 1C. Generally, the enrichable crosslinkers are more hydrophobic with larger structural space than the chain molecules, such as alkyne (azide)-A-DSBSO^[20]^ or CXL^[24]^. However, the crosslinker BDP-NHP^[17]^ (sometimes called PIR) with larger molecular size is not consistent the rule because of the better water solubility, which can ben interpreted as its internal polypeptide skeleton. The relatively large molecular length makes BDP-NHP more applicable to interaction studies than to conformation studies of protein complexes. Excitingly, the estimated values for Phox^[16]^, DSSO^[25]^ and BSP occupy a better performance with excellent water solubility and compact structural space. For Phox, the negative charges on phosphonic acid handle will preclude it from entering intact cells. And a high background of regular peptides will definitely inference DSSO in-vivo crosslinking. Therefore, the matched compact molecular size and better water solubility render BSP better membrane permeable capability, efficiently studying the conformation and interaction of protein complexes in living cells.

**Fig. 1.**
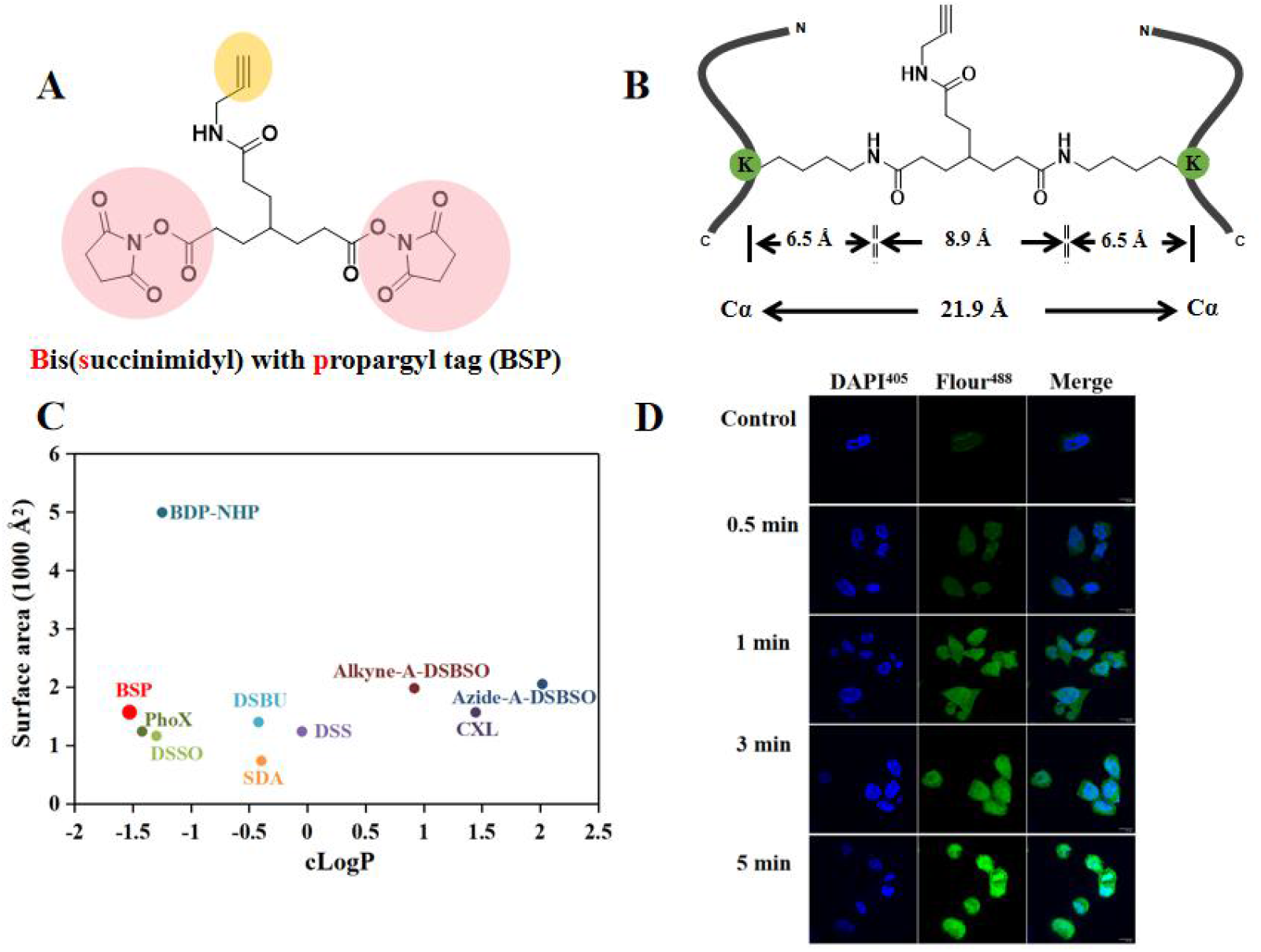
Chemical structure of BSP (A) and maximal distance constraint which is calculated by setting all interventing dihedral angles to 180° (B). Comparison on the amphiapathy and molecular size of BSP with the commonly reported crosslinkers containing NHS-ester group. cLogP value (horizontal axis) is a measure for the hydrophilicity and hydrophobicity of crosslinkers, which is obtained by ChemDraw Professional 17.0. The smaller the value, the better the hydrophilicity of the crosslinker. Surface area (vertical axis) calculations are accomplished by pyMol software (function ‘get_area’) to investigate the structural space of crosslinkers (C). Fluorescence confocal imaging for in-vivo crosslinking Bel7402 cell with BSP at differernt time (D).

### High reaction efficiency targeting amino of lysine

In order to evaluate the reactivity and enrichment properties of BSP, a model peptide Ac-SAKAYEHR containing a single amino group on lysine side chain was crosslinked and analyzed by Maldi-Tof (Figure 2A) and Lumos Mass spectrometry (Figure 2B), respectively. The crosslinking reaction was conducted in 5 min, which the crosslinked peptide were readily observed. After crosslinking, the sample was then subjected to the click chemistry reaction by cycloaddition with the diazo biotin-alkyne (DBA) probe for downstream streptavidin binding. The DBA probe (SP) was widely used for alkynyl-azide click chemistry reaction for the reason that the azo group could be efficiently cleaved with sodium dithionite in a mild procedure. Strong cation exchange (SCX) was adopted for the removal of excess unreacted DBA reagents to avoiding the potential competitive capture. After captured by streptavidin agarose resin (Thermo Fisher Scientific) and extensive washing, the biotinylated peptides could be released by sodium dithionite, resulting in biotin moiety left on the eluted peptides. MS^2^ fragmentation of crosslinked peptide, biotinylated crosslinked peptide and peptide released from avidin resin were performed and the ms2 spectra were analysed manually, assisted by pLabel software for spectrum labeling, de novo sequencing, yielding more b & y ions, which is helpful for the identification of the cross-linked peptide sequence. These results demonstrated BSP better performance of crosslinking, biotinylation through click chemistry and enrichment by streptavidin resin, allowing the possibility and availability of crosslinking with more complex biological samples.

**Fig. 2.**
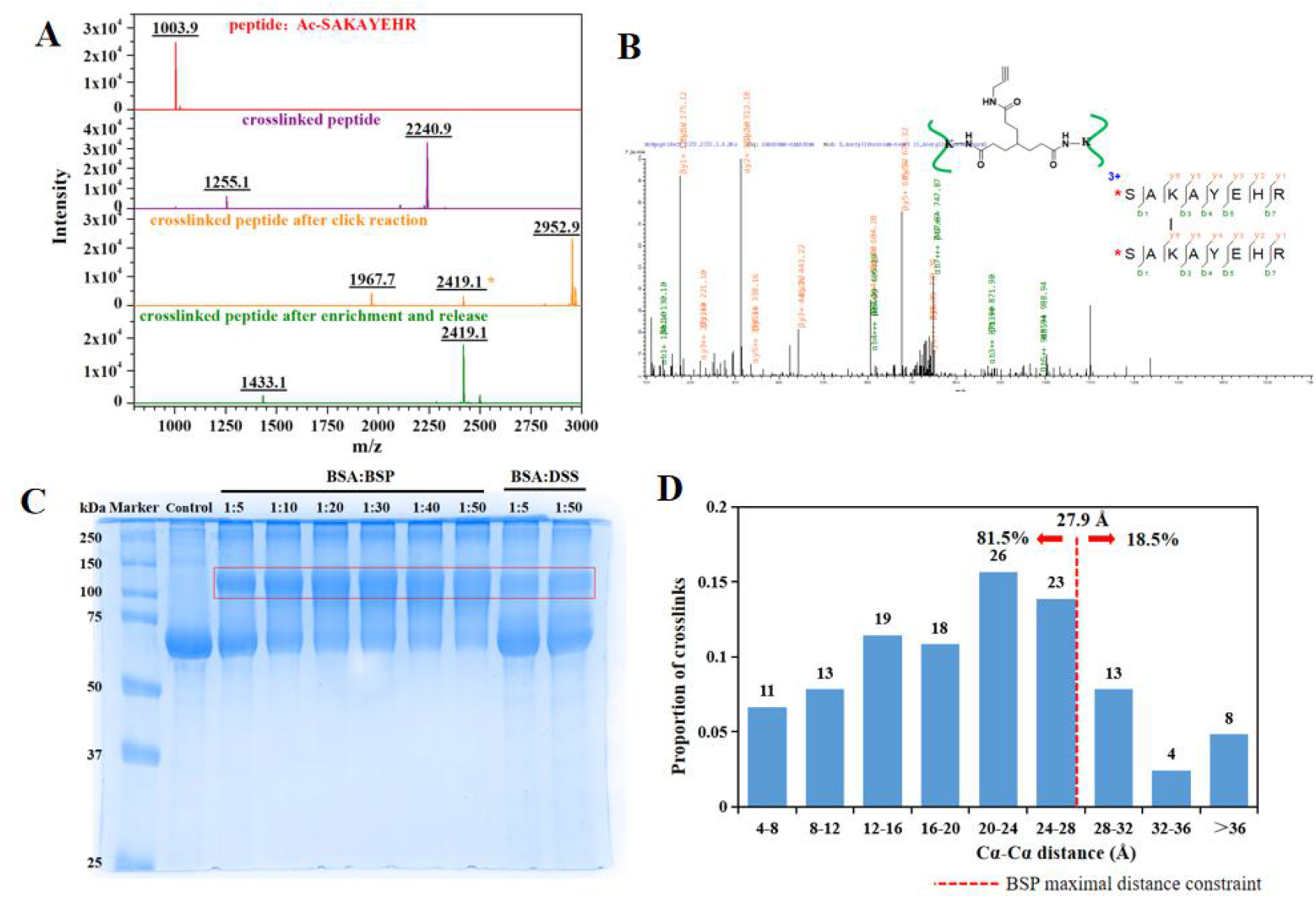
BSP crosslinking with model peptide (A), MS2 spectra of cross linked peptide (B), crosslinking with BSA (C) and the distribution of Cα–Cα distances of crosslinked residues of BSA (D).

To further assess the applicability of BSP in a protein environment, bovine serum albumin (BSA, a molecular weight of ∼ 66 kDa) was selected as model protein, which was widely used for testing crosslinking performance by SDS-PAGE analysis. We tested different molar ratio of BSA to BSP and benchmarked it against the widely used crosslinker disuccinimidyl suberate (DSS). Expected dimer bands were clearly detected on SDS-PAGE compared with the blank control and the single BSA band gradually weakened with the increase of BSP. Excitingly, the crosslinking performance of BSP was obviously superior to that of DSS from the dimer bands on SDS-PAGE (Figure 2C). That’s because our BSP is far more hydrophilic than DSS and no reactive sterical hindrance is introduced by the enrichable propargyl tag.

For BSA crosslinked sample, E-value ≤ 1e-4, PSM ≥ 4 were adoption to futher enhance the Identification accuracy, we identified 167 crosslinked peptides from click reaction workflow, demonstrating the excellent crosslinking efficiency of BSP. We further matched the 167 site pairs obtained from BSP with the BSA crystal structure to test the structural compatibility. The solvent accessible surface distance (SASD) of BSA was calculated using Xwalk. Because the crosslink site pairs (either intra- or inter-molecular) could not be determined based on the sequences of the crosslinked peptides, we then calculated all the possible combinations and selected the shorter Cα–Cα distance. The maximum Ca-Ca distance of BSP crosslinks is calculated to be 27.9 Å (PyMOL), and 78% (130 out of 167) of the identified cross-linked K–K pairs fall within the limit, showing a higher structural compatibility rate (Figure 2D). The over-length crosslinks can be attributed to the flexibility and dynamic of BSA, and the existence of higher protein aggregates.

### Better cell membrane permeability for in-vivo crosslinking

Finally, inspired by the good amphipathy, size and reactivity of BSP, in-vivo crosslinking performance was investigated by fluorescent confocal mocroscopy though click chemistry reaction with FITC-N3 in cells (Figure 1D). With the prolongation of crosslinking time, fluorescence intensity of the whole cells gradually increased, and reached the maximum fluorescence intensity in 5 min, confirming the good membrane permeability of BSP to enable in-situ immobilization of the structure of protein complexes in living cells.

### Development of efficient enrichment method based on click chemistry

After harvesting the cells, BSP dissolved in PBS was added and crosslinked with cells for 5 min to minimize cell disturbance as possible. The crosslinked proteins with alkyne moiety were obtained after cell lysis. After digested with trypsin by two rounds, the crosslinked peptides will be introduced with a alkyne moiety which can be effectively and specifically conjugated with biotin handle by copper(I)-catalyzed azide−alkyne cycloaddition (CuAAC) reaction. Then the crosslinked peptides can be enriched utilizing the highly specific biotin−streptavidin system to greatly improve the sensitivity of mass spectrometry. Recently, a new method of click chemistry reaction that took place at the peptide level was developed, resulting in a 2-fold increase in the identifications than that of protein level, due to the potential steric hindrance in proteins^[23]^. After enriched by avidin resin and harsh washing, a mild elution was exploited to release the crosslinked peptide by reduction cleavage that the azo group in DBA could be efficiently cleaved in sodium dithionite buffer. It is worth mentioning that the excess unreacted DBA reagents after click chemistry reaction should be removed by SCX separation, given that it would compete with the biotinylated peptides for streptavidin binding and enrichment. The released crosslinked peptides were subjected to standard shotgun proteomics mass spectrometry workflows and the data were searched against BSA or UniprotKB human protein database by pLink 2.0 software using an FDR of 1% as a filter at PSM level. The workflow was summarized in Figure 3A.

**Fig. 3.**
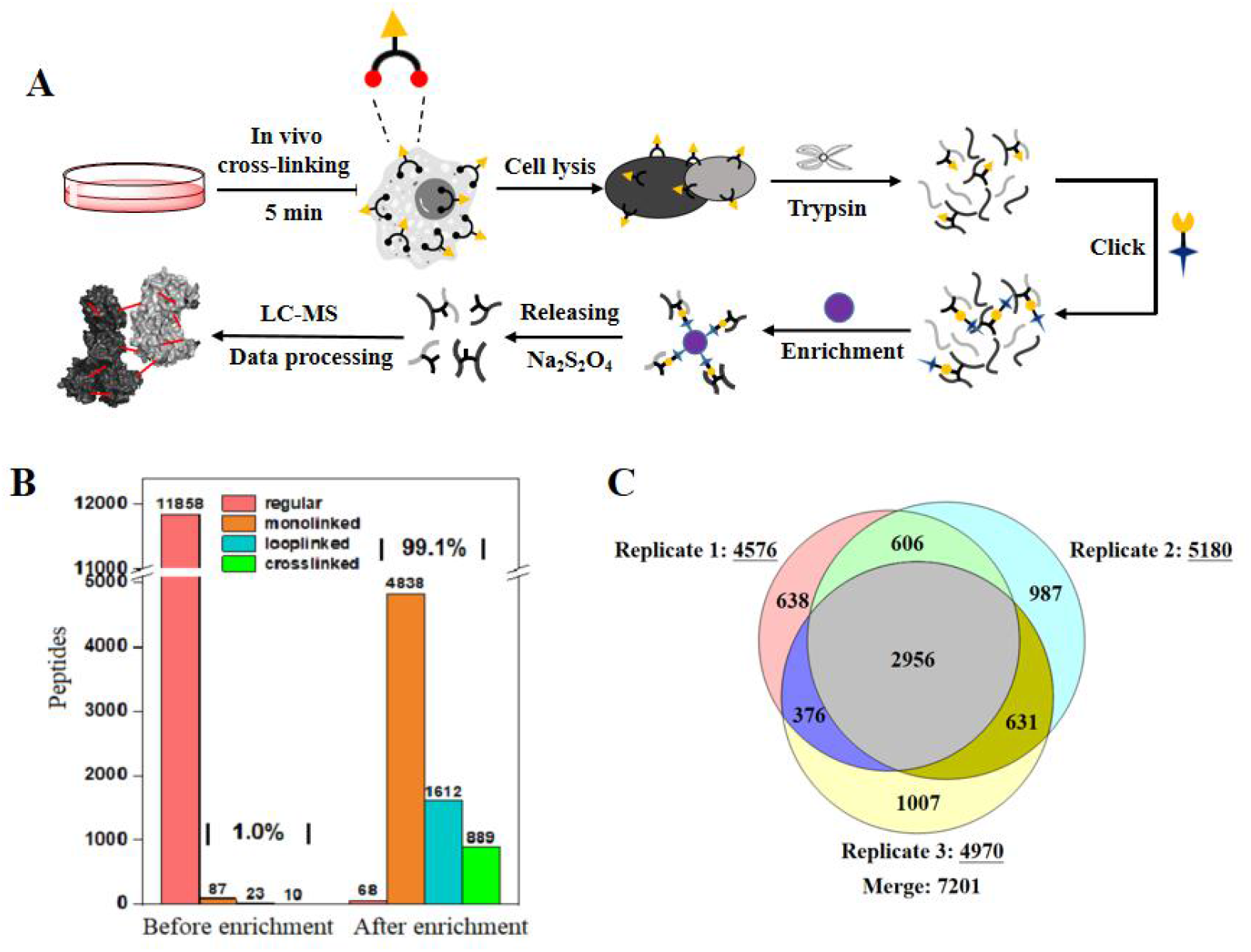
Flow diagrams of in vivo crosslinking (A), enrichment efficiency of crosslinked peptides based on click chemitry (B), venn diagrams of the identified crosslinks with 20 fractions derived from high pH C18 fractionation under 1% FDR control with three mass replicates (C).

To further evaluate the crosslinking performance of BSP and test the click chemistry approaches of conjugating with DBA probe in complex biological samples, in vivo crosslinking of intact Bel7402 cells was performed followed by downstream analysis by 150 min LC-MS acquisition without peptide fractionation. Before enriched by avidin resin, only 10 crosslinked peptides (accounts for 0.1%) were detected under the extreme background interference of the regular peptides (the faction was up to 99%), which demonstrated the necessity and importance of the enrichable crosslinker in CLMS. After enrichment, the regular peptides were decreased to a low percentage, leading to identification of more crosslinked peptides, which exhibited an excellent enrichment efficiency by biotin−avidin system. The detailed results of the click reaction workflow in technical triplicate were shown in venn diagram and Figure 3B.

### Profiling protein complexes by in-vivo crosslinking of living cells

In order to maximize the yield of the crosslinked peptides and gain more information of the protein structures and PPIs in situ, The sample of in vivo crosslinking of Bel7402 cells was further subjucted to the high pH Reverse-Phase C18 fractionation to afford 20 fractions. After 85 min gradient of LC-MS acquisition in technical triplicate, the raw data were searched by pLink and finally 7201 unique crosslinks could be identified at an FDR of 1%, 4898 (71.8%) of the cross-links were assigned as intraprotein, while 1922 (28.2%) were interprotein links. All cross-link data and protein abundance level data have been uploaded into XLinkDB and are accessible at the following URL…Our identification in human cells are better than those reported by other groups, due to the highly efficient enrichment ability of the crosslinked peptides. Protein interaction network generated exclusively from interprotein links are shown in Figure 4. Network consists of 2431 nodes representing proteins connected by 5152 edges representing interprotein cross-links. The interaction network was divided into several groups, according to the function of the protein and its location in the cell. Functional divisions include HSP, Oxidoreducatase, Histone, DNA binding, RNA binding ATP bingding proteins Zinc finger proteins and some low abundance proteins such as Transcription factor, nuclear pore complexes and kinase binding proteins. Protein location include protein complexes of various suborganelles, including ribosomal, mitochondrial, golgi, nucler pore and transmembrane proteins which demonstrated attractive ability of BSP for the large scale studies of the PPIS in cells. To better understand the significance of the interaction proteins, we used DAVID (Database for Annotation, Visualization and Integrated Discovery) database to perform the gene onthology term (GO term) analyses (Top ten proteins by P value in Fig). Detailed in Figure 5. The cellular component included nucleus, cytosol, membrane, mitochondrion and so forth. The molecular function annotation analysis includes RNA binding, ATP binding, kinase, DNA binding and so on, which were consistent with protein localization analysis in Fig. The interacting proteins we identified involved in a variety of celular biological processes, such as RNA processing, protein cotranslations, cell adhesion, cell division and so on, which were also related to protein functions. We further investigated the abundance of the crosslinked proteins (intra- and inter proteins) and compared the results with widely used crosslinker DSSO and PIR. Box plots of the abundance of identified proteins derived from proteome-wide crosslinked human cells by BSP, indicating our identifications were able to discover more proteins with low abundance than that of DSSO and PIR crosslinker, showing an attractive crosslinking ability of BSP.

**Fig. 4.**
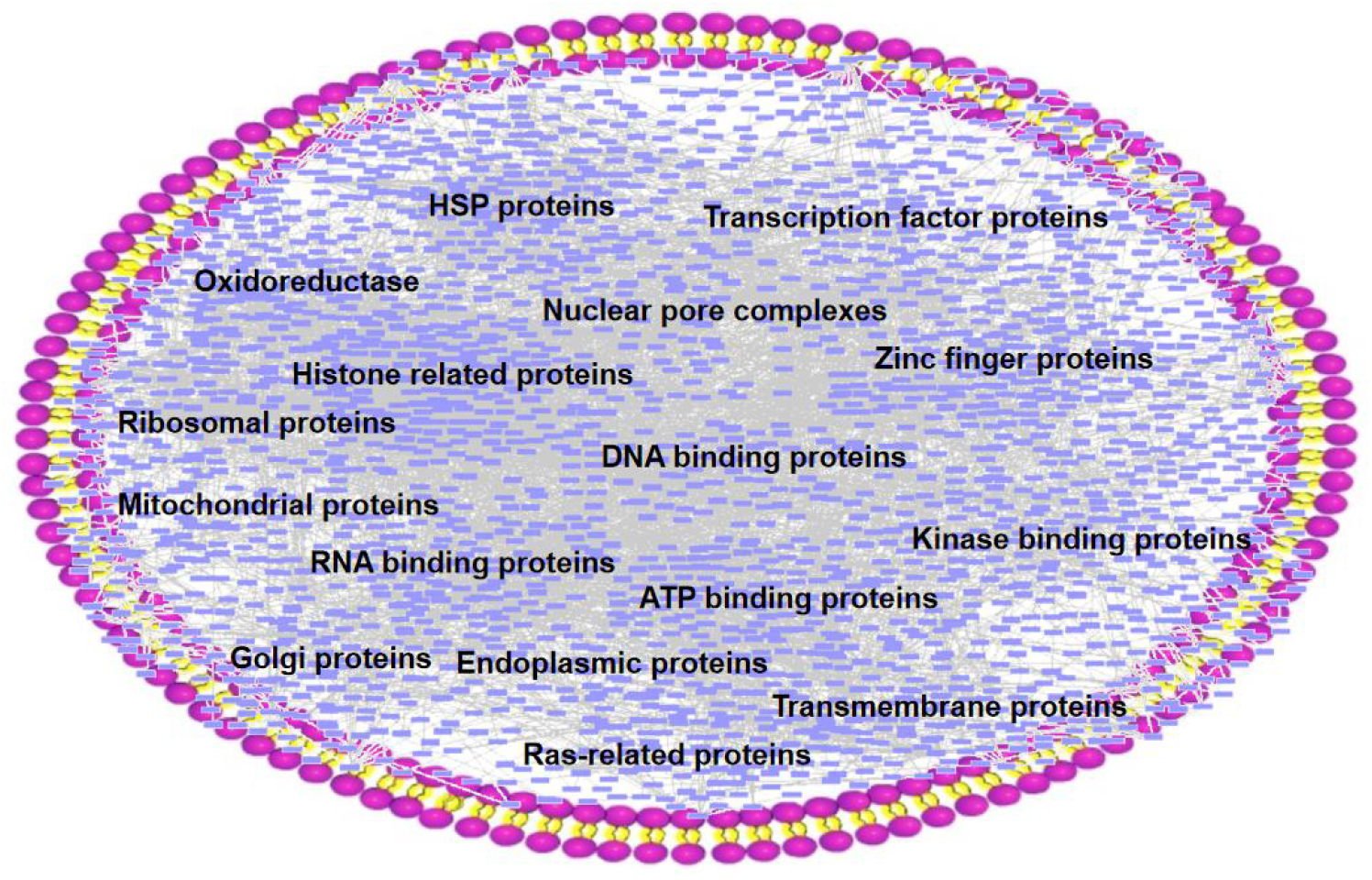
Protein interaction network generated exclusively from cross-linking results. Network consists of 2431 nodes representing proteins connected by 5152 edges representing interprotein cross-links. The interaction network was divided into several regions according to the localization and function of intracellular proteins.

**Fig. 5.**
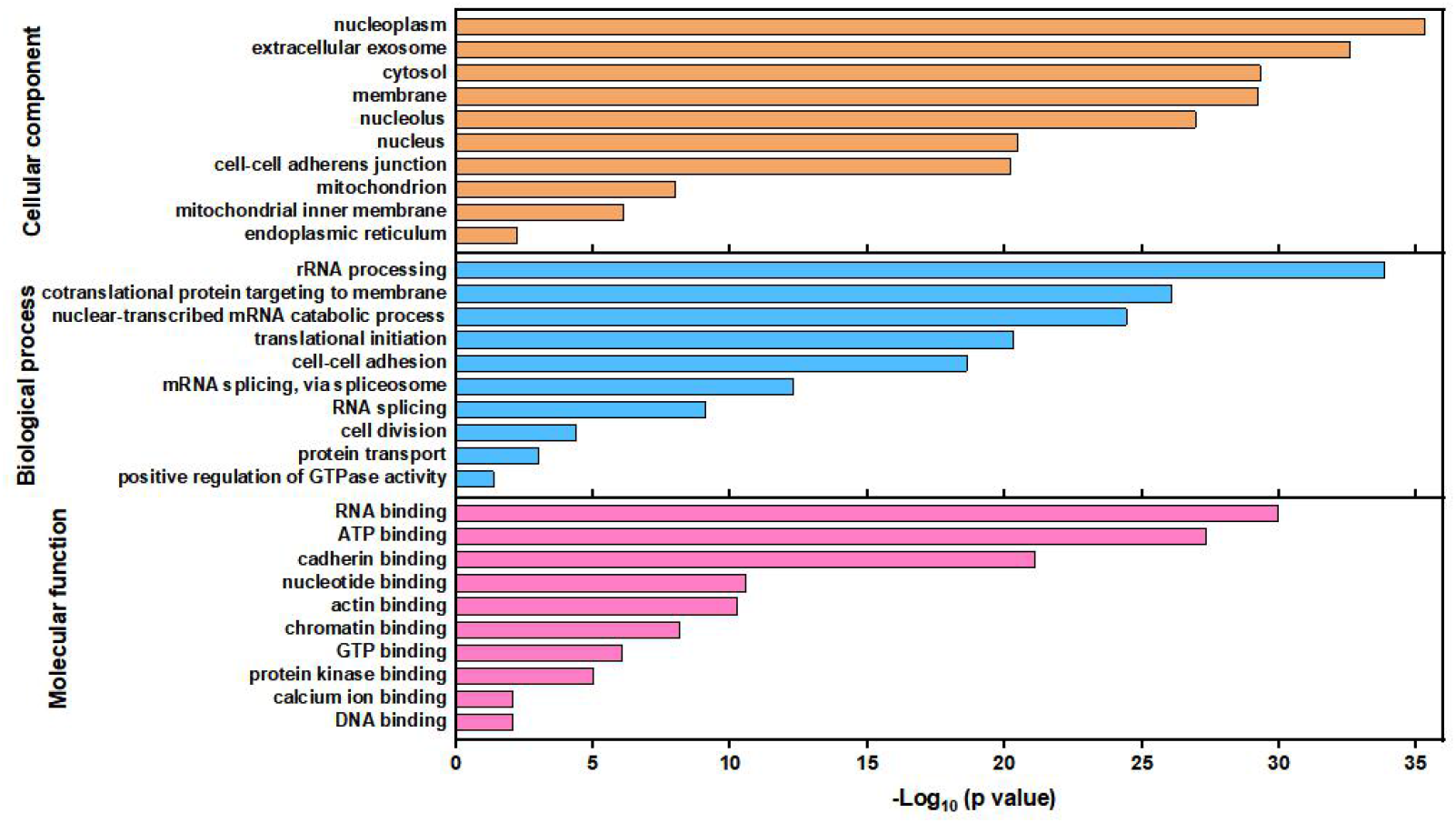
GO term analysis of cellular component (GOTERM_CC), biological processes (GOTERM_BP) and molecular function (GOTERM_MF) using DAVID program.

## Conclusion

Here we report the development and characterization of a new compact alkynyl-enrichable crosslinker, which has features of better membrane penetration, higher reactivity targeting amino of lysine side chain and excellent enrichable ability from complex environment. Thanks to the new in vivo CXMS pipeline based on click chemistry enrichment, we studied the protein complexes of human cells Bel7402, and finally identified 6820 crosslinks, including 4898 intraprotein links and 1922 interprotein links, demonstrating the promising prospect of BSP for in vivo studies. This is also the first time to realize the in-vivo crosslinking with a non-cleavable cross-linker for homo species cell. The development of this toolkit will facilitate the understanding of protein complexes in living cells.

## Supporting information

synthesized (detailed in supplementary material) consisting of two lysine-targeting reactive groups

## Acknowledgement

This work was supported by National Key Research and Development Program of China (2016YFA0501401).

## References

[1] Iacobucci, C.; Götze, M.; Sinz, A. Curr. Opin. Biotechnol. 2020, 63, 48.

[2] Chavez, J. D.; Bruce, J. E. Curr. Opin. Chem. Biol. 2019, 48, 8.

[3] Yu, C.; Huang, L. Anal. Chem. 2018, 90, 144.

[4] Schneider, M.; Belsom, A.; Rappsilber, J. Trends Biochem. Sci. 2018, 43, 157.

[5] Tan, D. Ph.D. Dissertation, National Institute of Biological Sciences, Beijing, 2014 (in Chinese).

[6] Chu, F.; Thornton, D. T.; Nguyen, H. T. Methods 2018, 144, 53.

[7] Steigenberger, B.; Albanese, P.; Heck, A. J. R.; Scheltema, R. A. J. Am. Soc. Mass. Spectrom. 2020, 31, 196.

[8] Zybailov, B. L.; Glazko, G. V.; Jaiswal, M.; Raney, K. D. J. Proteomics Bioinform. 2013, 6, 001.

[9] Leitner, A.; Faini, M.; Stengel, F.; Aebersold, R. Trends Biochem. Sci. 2016, 41, 20.

[10] Liu, F.; Lossl, P.; Scheltema, R.; Viner, R.; Heck, A. J. R. Nat. Commun. 2017, 8, 15473.

[11] Kastritis, P. L.; O’Reilly, F. J.; Bock, T.; Li, Y.; Rogon, M. Z.; Buczak, K.; Romanov, N.; Betts, M. J.; Bui, K. H.; Hagen, W. J.; Hennrich, M. L.; Mackmull, M. T.; Rappsilber, J.; Russell, R. B.; Bork, P.; Beck, M.; Gavin, A. C. Mol. Syst. Biol. 2017, 13, 936.

[12] Schweppe, D. K.; Chavez, J. D.; Lee, C. F.; Caudal, A.; Kruse, S. E.; Stuppard, R.; Marcinek, D. J.; Shadel, G. S.; Tian, R.; Bruce, J. E. Proc. Natl. Acad. Sci. U. S. A. 2017, 114, 1732.

[13] Zhong, X.; Wu, X.; Schweppe, D. K.; Chavez, J. D.; Mathay, M.; Eng, J. K.; Keller, A.; Bruce, J. E. J. Am. Soc. Mass. Spectrom. 2019, 31, 190.

[14] Chavez, J. D.; Schweppe, D. K.; Eng, J. K.; Bruce, J. E. Cell Chem. Biol. 2016, 23, 716.

[15] Chavez, J. D.; Lee, C. F.; Caudal, A.; Keller, A.; Tian, R.; Bruce, J. E. Cell Syst. 2018, 6, 136.

[16] Liu, S.; Yu, F.; Hu, Q.; Wang, T.; Yu, L.; Du, S.; Yu, W.; Li, N. J. Proteome Res. 2018, 17, 3195.

[17] Chavez, D. J.; Keller, A.; Zhou, B.; Tian, R.; Bruce E. J.; Cell Reports, 2019, 29, 2371–2383.

[18] Tan, D.; Li, Q.; Zhang, M.-J.; Liu, C.; Ma, C.; Zhang, P.; Ding, Y.-H.; Fan, S.-B.; Tao, L.; Yang, B.; Li, X.; Ma, S.; Liu, J.; Feng, B.; Liu, X.; Wang, H.-W.; He, S.-M.; Gao, N.; Ye, K.; Dong, M.-Q.; Lei, X. eLife 2016, 5.

[19] Steigenberger, B.; Pieters, R. J.; Heck, A. J. R.; Scheltema, R. A. ACS Cent. Sci. 2019, 5, 1514.

[20] Kaake, R. M.; Wang, X.; Burke, A.; Yu, C.; Kandur, W.; Yang, Y.; Novtisky, E. J.; Second, T.; Duan, J.; Kao, A.; Guan, S.; Vellucci, D.; Rychnovsky, S. D.; Huang, L. Mol. Cell. Proteomics 2014, 13, 3533.

[21] Chavez, J. D.; Weisbrod, C. R.; Zheng, C.; Eng, J. K.; Bruce, J. E. Mol. Cell. Proteomics 2013, 12, 1451.

[22] Luo, J.; Fishburn, J.; Hahn, S.; Ranish, J. Mol. Cell. Proteomics 2012, 11, M111.008318.

[23] Sun N.; Wang, Y.; Wang, J.; Sun W.; Yang J.; Liu N.; Anal. Chem., 2020, 92, 8292–8297.

[24] Sohn, C. H.; Agnew, H. D.; Lee, J. E.; Sweredoski, M. J.; Graham, R. L. J.; Smith, G. T.; Hess, S.; Czerwieniec, G.; Loo, J. A.; Heath, J. R.; Deshaies, R. J.; Beauchamp, J. L. Anal. Chem. 2012, 84, 2662.

[25] Ser, Z.; Cifani, P.; Kentsis, A. J. Proteome Res. 2019, 18, 2545.

