## Supplementary material for "Development of a compact alkynyl-enrichable crosslinker for in-depth in-vivo crosslinking analysis": synthesized (detailed in supplementary material) consisting of two lysine-targeting reactive groups

### *EXPERIMENTAL PROCEDURES*

#### *Instruments and methods for the seperation and characterization of crosslinker*

The process of reactions and purity of products were monitored by analytical HPLC from HITACHI chromaster (Japan) equipped with a DAD UV detector using C18 column (3 µm, 100 Å, 4.6 × 250 mm i.d.).

Semi-preparative RP-HPLC purifications were performed on DAD50 system from Hanbon (China) installed with a UV-visible 2000 detector using C18 column (10 µm, 100 Å, 50 × 250 mm i.d.). The freeze-drying of crosslinker purified by DAD50 HPLC system was carried out on a lyophilizer from Tianrey (China) at 30 °C for 24 h. $^{1}H$ and $^{13}C$ NMR spectra were recorded on a Bruker AVANCE II 400 MHz spectrometer using $CDCl_3$ or DMSO-d6 as the solvent and TMS as the internal standard. High-resolution mass spectra (HRMS) were recorded on an LTQ Orbitrap Velos (Thermo Fisher Scientific, Bremen, Germany).

#### *Synthesis of BSP*

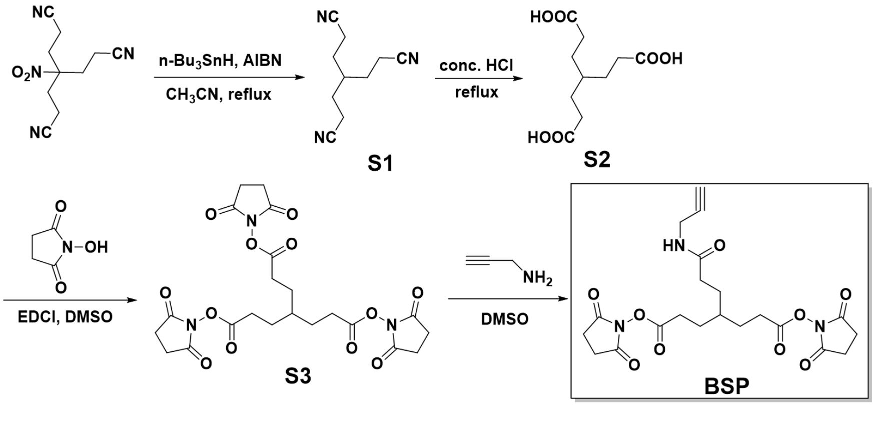


**Figure S1. Synthesis route of BSP**

***4-(2-cyanoethyl)heptanedinitrile (trisnitrile, S1)***

A mixture of tris(2-cyanoethyl)nitromethane (9.2 g, 40 mmol), tri-n-butyltin hydride (14.0 g, 48 mmol) and AIBN (2.36 g, 14.4 mmol) in 300 mL HPLC-grade $CH_3CN$ was refluxed at 88 °C under $N_2$ atmosphere for 12 h. After being cooled to the room temperature, the organic solvent was removed under reduced pressure and the crude product was recrystallized twice with 50 ml methanol. The precipitate was washed with little cool methanol and dried in vacuo for 24 h to afford a white solid compound S1 (6.31 g, yield 90.1%). $^{1}H$ NMR (400 MHz, $CDCl_3$, ppm) δ 2.45 (t, 6H, *J* = 7.2 Hz), 1.83-1.73 (m, 7H). LTQ Mass: Exact mass 175.1109 for C10H13N3, found 198.1230 $[M+Na]^{+}$.

***4-(2-carboxyethyl)heptanedioic acid (tris-carboxylic acid, S2)***

Compound S1 (6.1 g, 35 mmol) was refluxed in conc. HCl (40 ml) for 1 h and then cooled to

0 °C. The precipitate was filtered and washed with little cooled water, dried in vacuo for 24 h to afford a white solid compound S2 (7.2 g, yield 88.7%). $^{1}H$ NMR (400 MHz, DMSO-d6, ppm) δ 12.03 (m, 3H), 2.20 (t, 6H, $J$ = 7.6 Hz), 1.48-1.43 (m, 6H), 1.35-1.31 (m, 1H). LTQ Mass: Exact mass 232.0947 for C10H16N6, found 250.1298 $[M+H_2O]^+$.

***Tris-succinimide ester (S3)***

Compound S2 (1.86 g, 8 mmol), EDCI (4.14g, 36 mmol), NHS (6.9g, 36 mmol) were dissolved in 100 ml anhydrous $CH_2Cl_2$ and stirred at room temperature for 24 h. After being detected for completion by analytical HPLC, the solution was pooled into 100 ml deionized $H_2O$ and washed with $H_2O$ for three times. The organic layer was collected, dried over anhydrous $Na_2SO_4$ and removed under vacuum to afford a white solid compound S3 (3.98 g, yield 95.2%). Purity 99%, retention time at 28 min detected by analytical HPLC (A: $H_2O$, B: $CH_3CN$, gradient method was as follows: B from 5% to 50% over 40 min, 80% B over 10 min, 5% B over 10min at a flow rate of 1 mL/min monitored by UV wavelength of 200 nm). $^{1}H$ NMR (400 MHz, DMSO-d6, ppm) δ 2.81 (s, 12H), 2.71 (t, 6H, $J$ = 7.6 Hz), 1.70-1.65 (m, 6H), 1.62-1.57 (m, 1H); $^{13}C$ NMR (400 MHz, DMSO-d6, ppm) δ 170.7, 169.5, 35.5, 28.0, 27.3, 25.9. LTQ Mass: Exact mass 523.1438 for C22H25N3O12, found 524.0555 $[M+H]^+$.

***Trifunctional crosslinker (BSP)***

Compound S2 (1.05 g, 2 mmol) was dissolved in 30 ml DMSO and propargyl amine (110 mg, 2 mmol) in 1 ml DMSO was added dropwise to the solution. After being stirred at room temperature for 5 min and detected for completion by analytical HPLC, the mixture was purified by semi-preparative RP-HPLC (A: 0.1% TFA/H2O, B: $CH_3CN$, gradient method was as followed: B from 20% to 50% over 30 min at a flow rate of 50 mL/min monitored by UV wavelength of 200 nm). The product-containing fractions (retention time, 22.0 - 26.0 min) were lyophilized at 30 °C for 24 h to afford the targeted trifunctional crosslinker BSP as a light yellow sticky oil. Purity 99%, retention time at 26 min detected by analytical HPLC (A: 0.1% TFA/$H_2O$, B: $CH_3CN$, gradient method was as followed: B from 5% to 50% over 40 min, 80% B over 10 min, 5% B over 10min at a flow rate of 1 mL/min monitored by UV wavelength of 200 nm). $^{1}H$ NMR (400 MHz, DMSO-d6, ppm) δ 8.29-8.27 (m, 1H), 3.84 (t, 2H, $J$ = 2.6 Hz), 3.07 (s, 1H), 2.81 (s, 8H), 2.69 (t, 4H, $J$ = 7.8 Hz), 2.11 (t, 2H, $J$ = 7.6 Hz), 1.63-1.60 (m, 4H), 155-1.49 (m, 3H); $^{13}C$ NMR (400 MHz, DMSO-d6, ppm) δ 172.3, 170.7, 81.7, 73.3, 35.8, 32.5, 28.3, 28.1, 27.6, 25.9, 25.6. LTQ Mass: Exact mass 463.1591 for C21H25N3O9, found 464.2333 $[M+H]^+$.

***Compound Characterizations***

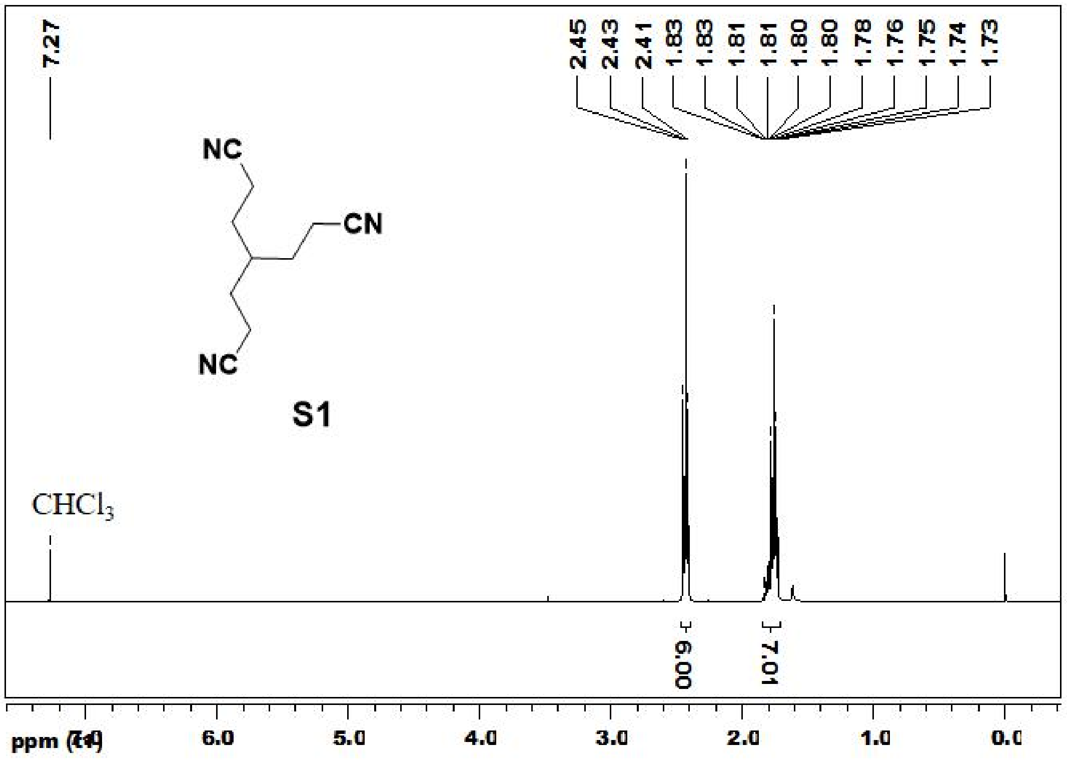


[1]H-NMR of compound S1

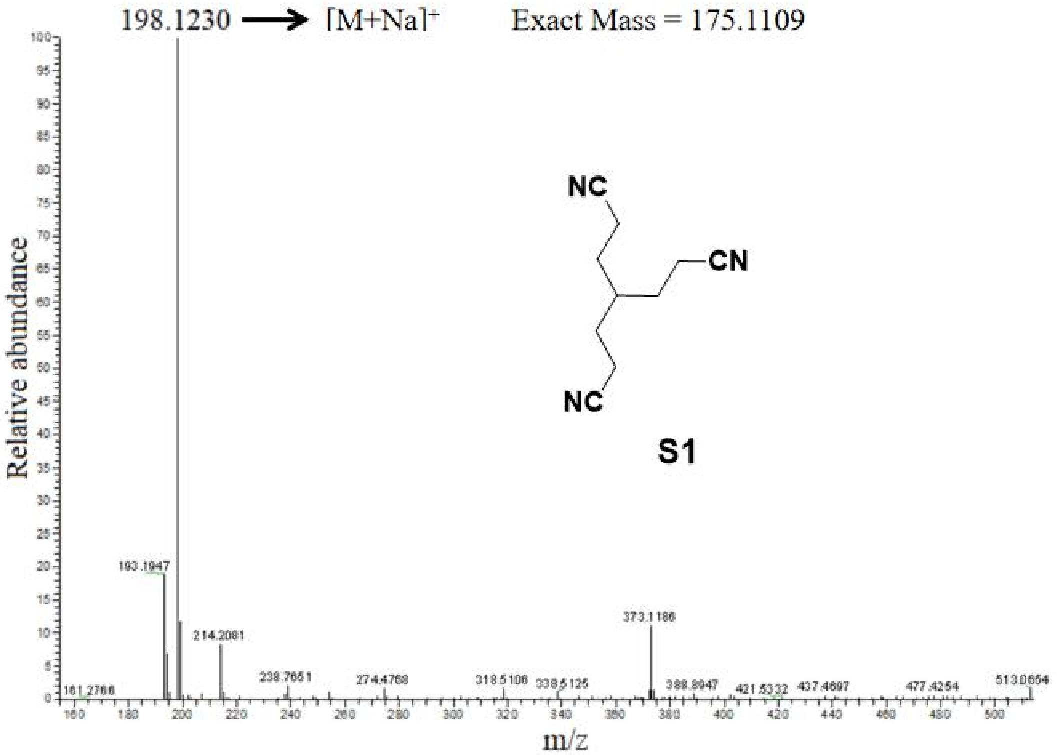


LTQ-Mass of compound S1

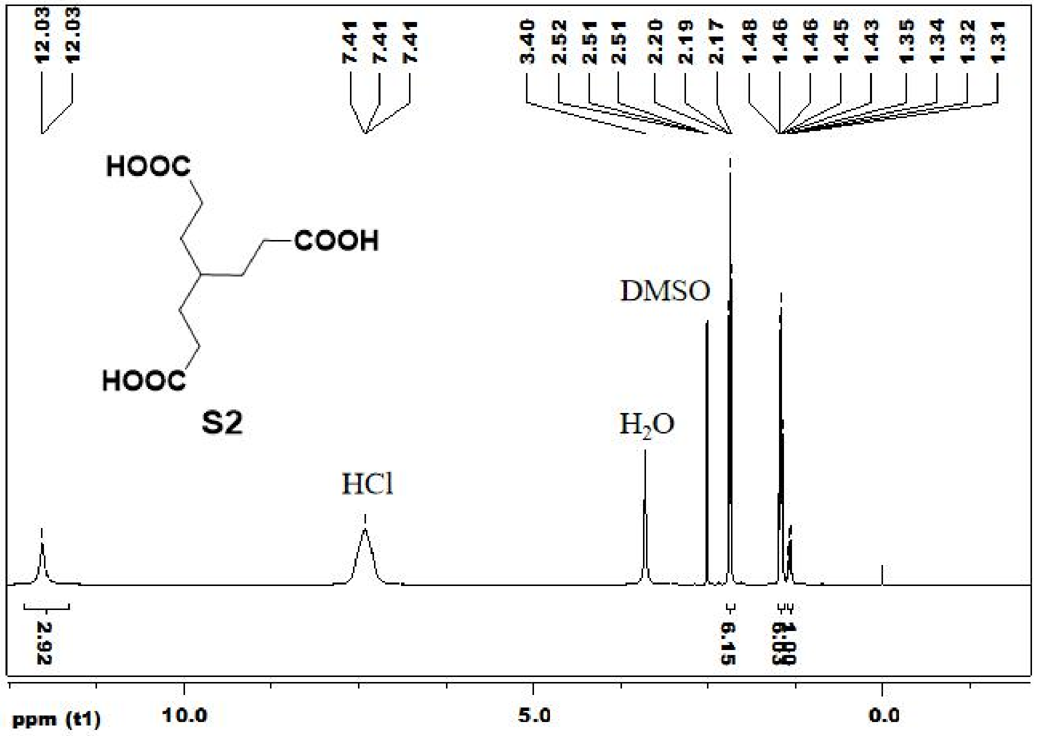


[1]H-NMR of compound S2

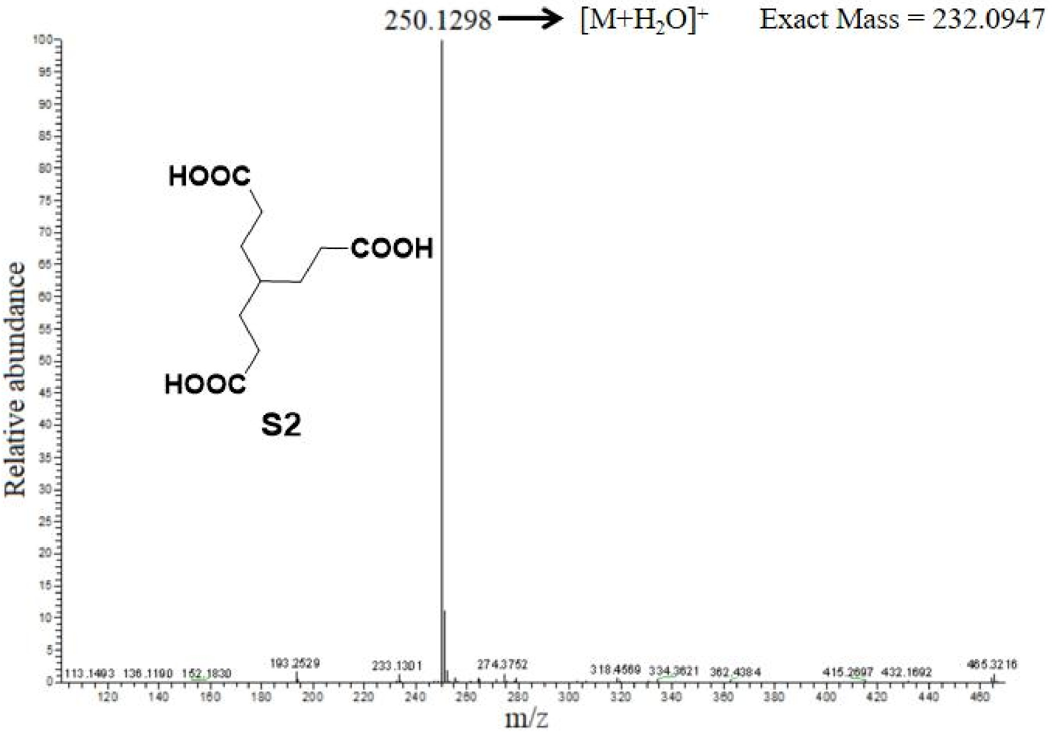


LTQ-Mass of compound S2

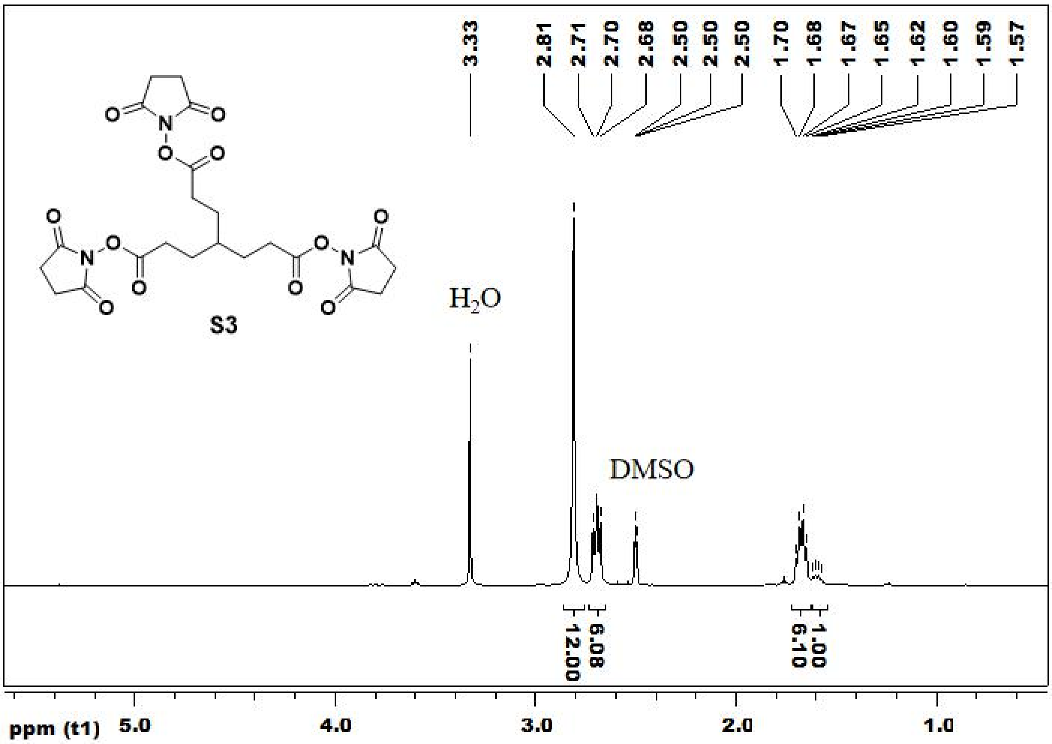


[1]H-NMR of compound S3

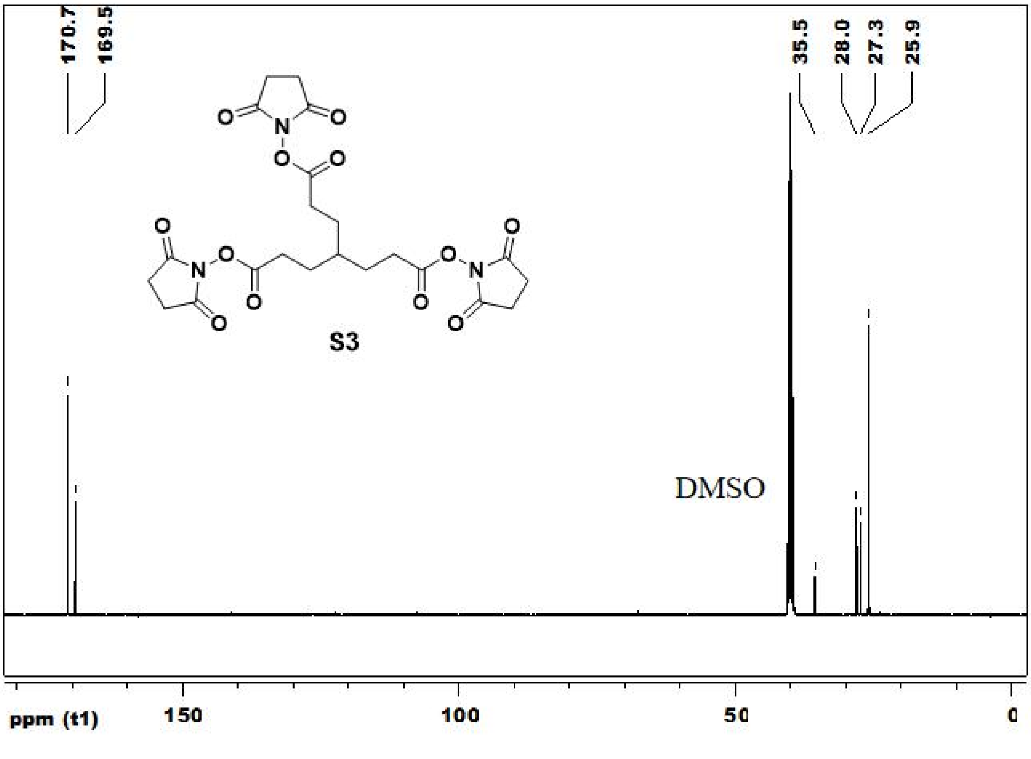


[13]C-NMR of compound S3

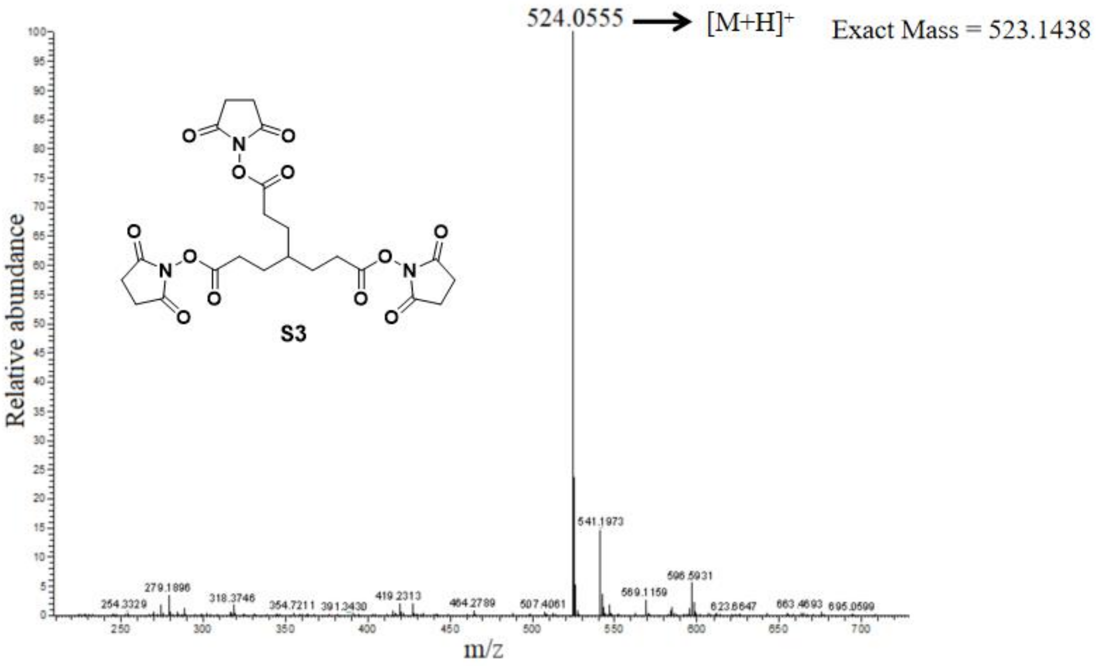


LTQ-Mass of compound S3

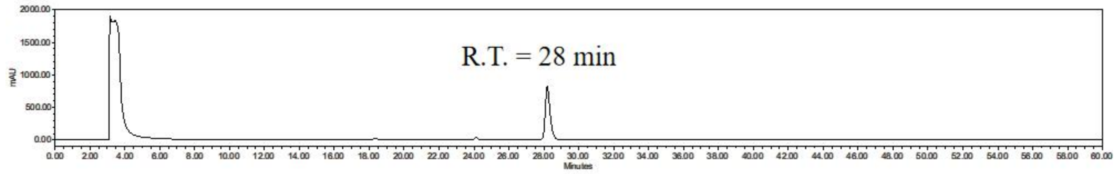


HPLC retention of compound S3

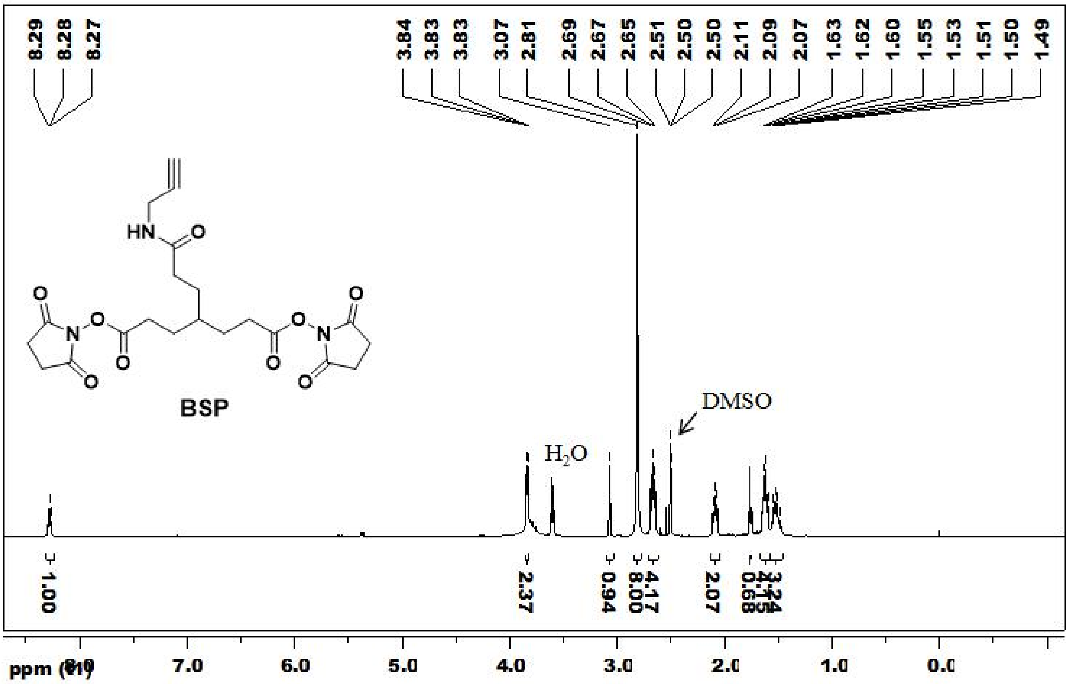


[1]H-NMR of compound BSP

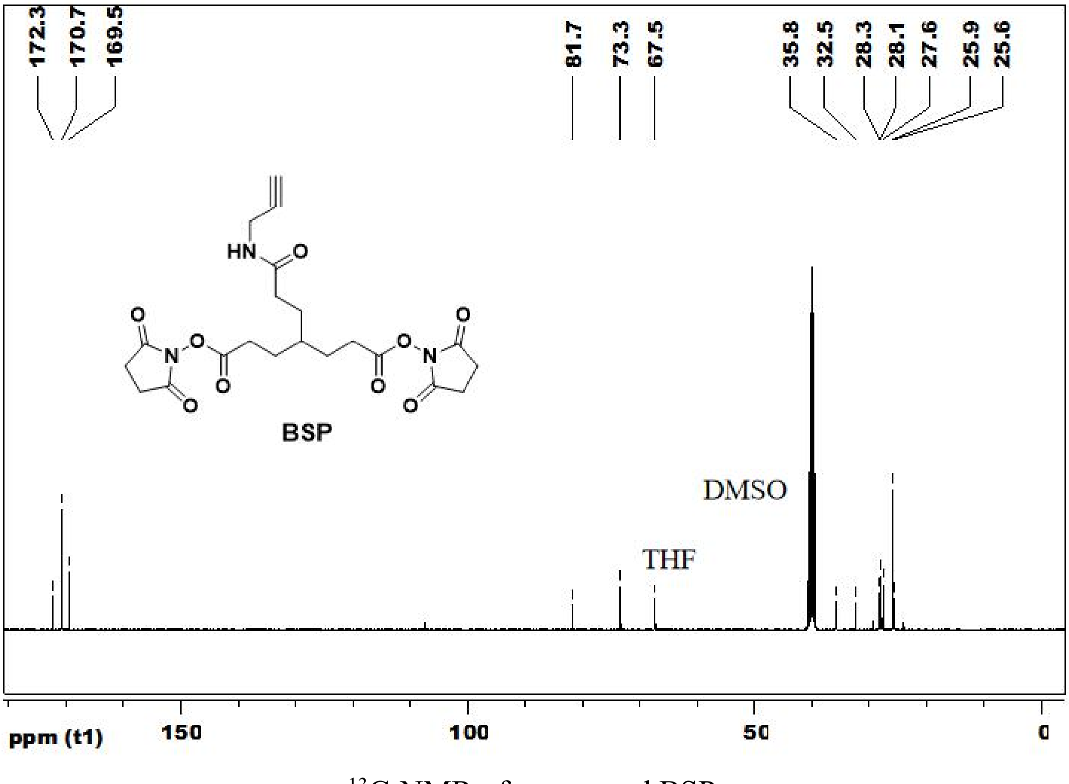


[13]C-NMR of compound BSP

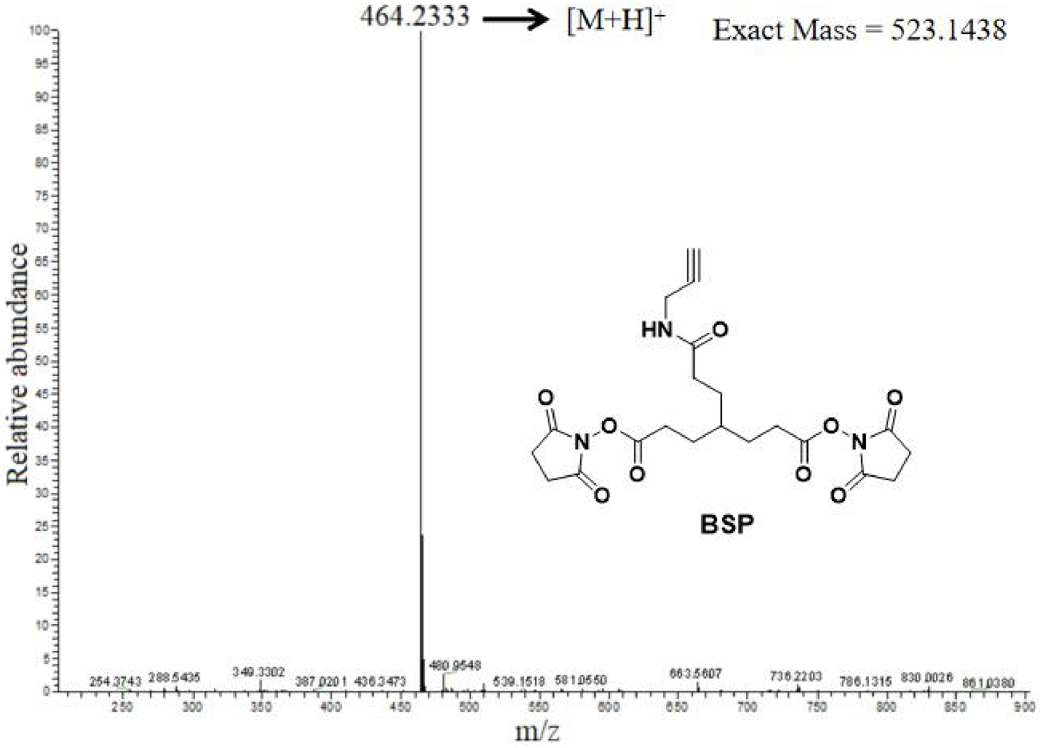


LTQ-Mass of compound BSP

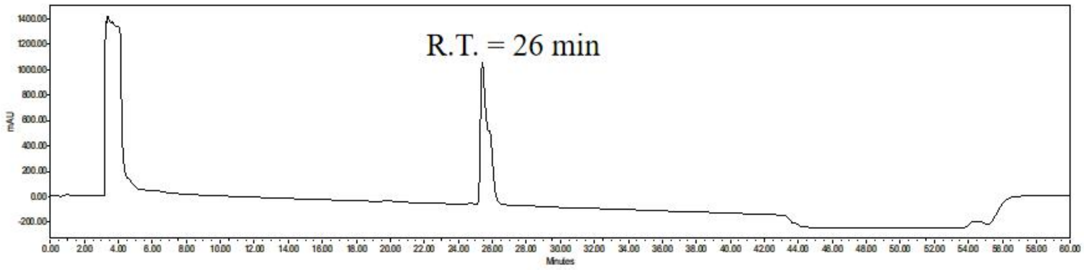


HPLC retention of compound BSP
